# An Augmented Reality Visor for Intraoperative Visualization, Guidance and Temperature Monitoring using Fluorescence

**DOI:** 10.1101/2024.08.08.607031

**Authors:** Oscar Cipolato, Matthias Fauconneau, Paige J. LeValley, Robert Nißler, Benjamin Suter, Inge K. Herrmann

## Abstract

Fluorescence-guided surgical techniques, including tumor resection and tissue soldering, are advancing the frontiers of surgical precision by offering enhanced control that minimizes tissue damage and improves recovery as well as outcomes. However, integrating the visualization of the fluorescent signal and temperature monitoring seamlessly into surgical workflows has not been fully realized and remains a challenge, thus limiting their effectiveness and wide-spread clinical adoption. To address this issue, we introduce an augmented reality (AR) visor designed to unite nanomaterial excitation along with fluorescence detection, and temperature monitoring during surgical procedures. The AR visor was evaluated using advanced fluorescent nanoparticles, such as indocyanine green-doped particles and single-walled carbon nanotubes. By consolidating fluorescence visualization, excitation monitoring, and precise temperature control into a single AR platform, we equip surgeons with a comprehensive view of both the surgical field and sub-surface conditions invisible to the naked eye. This integration notably improves the safety and efficacy of fluorescence-guided surgeries, as well as emerging technologies including laser tissue soldering, by ensuring the soldering temperature stays within therapeutic thresholds and the laser is accurately guided by real-time fluorescence signals. The presented technology not only enhances existing surgical techniques but also supports the development of new strategies and sensing technologies in areas where traditional methods fall short, marking significant progress in precision surgery, which could ultimately improve patient care.

## INTRODUCTION

Fluorescence-guided surgery has revolutionized certain surgical practices,(*1*–*4*) notably enhancing the surgeon’s ability to precisely identify and remove cancerous tissues while sparing vital structures.(*5*) This technique employs fluorescent dyes, typically excited and emitting in the near-infrared (NIR) region between 650 nm and 1350 nm known as biological window, to illuminate target areas and provide clear visualization of tumor margins within deep tissues. The use of NIR light is advantageous because it is less absorbed and scattered by bodily tissues, allowing deeper tissue penetration and making subsurface structures visible.(*6, 7*) Indocyanine green (ICG), a medically approved dye, is frequently utilized for this purpose.(*8, 9*) Advancements in the development of novel fluorescent materials are further enhancing the effectiveness of fluorescence-guided surgery. ICG-loaded nanoparticles (ICG NPs) are a notable example, utilizing a clinically validated dye for safe application. Not only do they enhance the stability of the dye, but they can also be engineered to optimize tumor uptake through the enhanced permeability and retention (EPR) effect.(*10, 11*) Alternatively, they can be modified to actively target tumors, increasing their precision.(*12, 13*) Carbon nanotubes (CNTs), especially fluorescent single-walled CNTs (SWCNTs), represent another promising class of nanomaterials rapidly advancing within fluorescence-guided surgical techniques.(*14*–*17*) SWCNTs can be engineered to enhance their imaging and thermal properties, as well as their emission wavelengths ranging from NIR-I to NIR-II without being prone to photo-bleaching, making them highly versatile for various medical applications including tumor hyperthermia.(*18*–*22*) Additionally, SWCNTs can be tailored to activate their fluorescence in response to specific conditions, such as the presence of tumor cells or other distinct biological environments, hence acting as chemical sensors.(*23*–*27*)

In addition to visualizing diseased tissue as well as vital structures, such as vessels and nerves, controlling temperature is critically important in various surgical procedures, especially in tumor treatment where deliberate increases in temperature are used for therapeutic effects, such as destroying malignant cells through controlled hyperthermia. Hyperthermia uses elevated temperatures (from 42 °C to above 50 °C) to target tumor cells without harming surrounding healthy tissue.(*28, 29*) However, precise temperature control is crucial to avoid complications like delayed healing or extended necrosis of healthy tissue. Similarly, when lasers illuminate vital structures during surgery, accurate temperature monitoring is essential to prevent inadvertent heating from excessive laser illumination, preserving healthy tissue integrity. While fluorescence-guided surgery improves tumor localization, it also necessitates tools for simultaneous temperature monitoring during surgical interventions.

The importance of fluorescence-guided operations are not limited to tumor visualization and hyperthermia applications. Laser tissue soldering is an emerging technique that represents an alternative to sutures for wound closure,(*30*–*36*) particularly in surgeries involving delicate tissues such as blood vessels,(*37*) nerves,(*38*) intestines,(*39, 40*) and dura mater.(*41*) Traditional suturing often leads to complications like slow healing, tears, leakages, and infections.(*42*–*45*) Laser tissue soldering utilizes lasers to heat biocompatible, protein-based solder pastes applied to wounds, aiming for effective closure with minimal tissue damage. This method is especially advantageous following tumor resection, where precise rejoining of tissues and watertight seals are necessary. However, the broader adoption of laser tissue soldering has been hindered by challenges in controlling the soldering temperature and the difficulty surgeons face in visualizing the laser spot size and position due to the need for protective goggles.(*31, 39*) Enhancing laser tissue soldering requires the integration of real-time information about the location and size of the laser, and the temperature of the tissue. Infrared thermometry, which monitors surface temperatures, and NIR cameras, provide crucial feedback during the procedure. Additionally, incorporating fluorescent additives into the solder paste—such as ICG for its heating properties or fluorescent nanothermometers for real-time temperature monitoring—enhances guidance, helping surgeons ensure the laser accurately targets the paste.

In the quest to advance surgical technology, augmented reality (AR) is transforming surgical procedures by superimposing critical, otherwise invisible information directly onto the surgeon’s visible field, enhancing spatial orientation and surgical guidance.(*46*–*48*) Unlike many AR systems that rely on additional monitors, which divert the surgeon’s attention from the operative field and increase stress, head-mounted AR displays integrate crucial data seamlessly into the surgeon’s line of sight.(*49*–*51*) This method is particularly effective in environments involving lasers, where safety goggles required for protection absorb visible light and reduce the clarity of displayed information. Additionally, AR visors that present information before the laser filter can display a broader range of colors, improving communication of diverse surgical data. By seamlessly integrating with laser tissue soldering and other fluorescence-guided techniques, AR not only addresses the challenges of managing complex arrays of surgical information but also promises substantial improvements in surgical outcomes. Importantly, this technology can be an affordable solution, making advanced surgical guidance accessible to a wider range of healthcare facilities.

In this work, we introduce a custom AR visor designed for fluorescence-guided surgeries that involve visualization of blood vessels and tumors, together with controlled heating, and laser tissue soldering, effectively integrated into the workflow of tumor removal. We demonstrate the visor’s capability to accurately locate and controllably heat fluorescently-labeled regions, as well as facilitate precise and safe soldering by providing real-time data on laser spot position and size, laser-paste co-localization and color-coded temperature information. Additionally, we showcase the visor’s utility in rapidly detecting leakages or blockages post-wound closure in fluid-containing structures like blood vessels. This is demonstrated using two different types of nanoparticles—ICG-loaded nanoparticles and SWCNTs— highlighting the versatility of our platform and its role in facilitating the adoption of novel technologies.

## RESULTS & DISCUSSION

### Design of the augmented reality visor

To advance image-guided surgical technology and give the surgeon capabilities beyond natural vision, we have developed an AR visor designed to enhance surgical precision through NIR fluorescence imaging and infrared temperature imaging, aimed at applications such as tumor resection and hyperthermia, wound closure via laser tissue soldering, and perfusion tests for leak detection and vascular identification (Figure 1a-b). The system utilizes a 750 nm laser with a maximum output of 2.5 W as the excitation source, selected for its dual capability to both excite a wide range of fluorophores and facilitate laser tissue soldering, providing a versatile tool in surgical settings. The laser illumination is delivered through an optical fiber with numerical aperture NA = 0.22, which is handheld by the user. The absence of collimating optics at the tip of the fiber allows the operator to straightforwardly modify the illuminated area and laser intensity by adjusting the distance between the fiber and the target.

**Figure 1:**
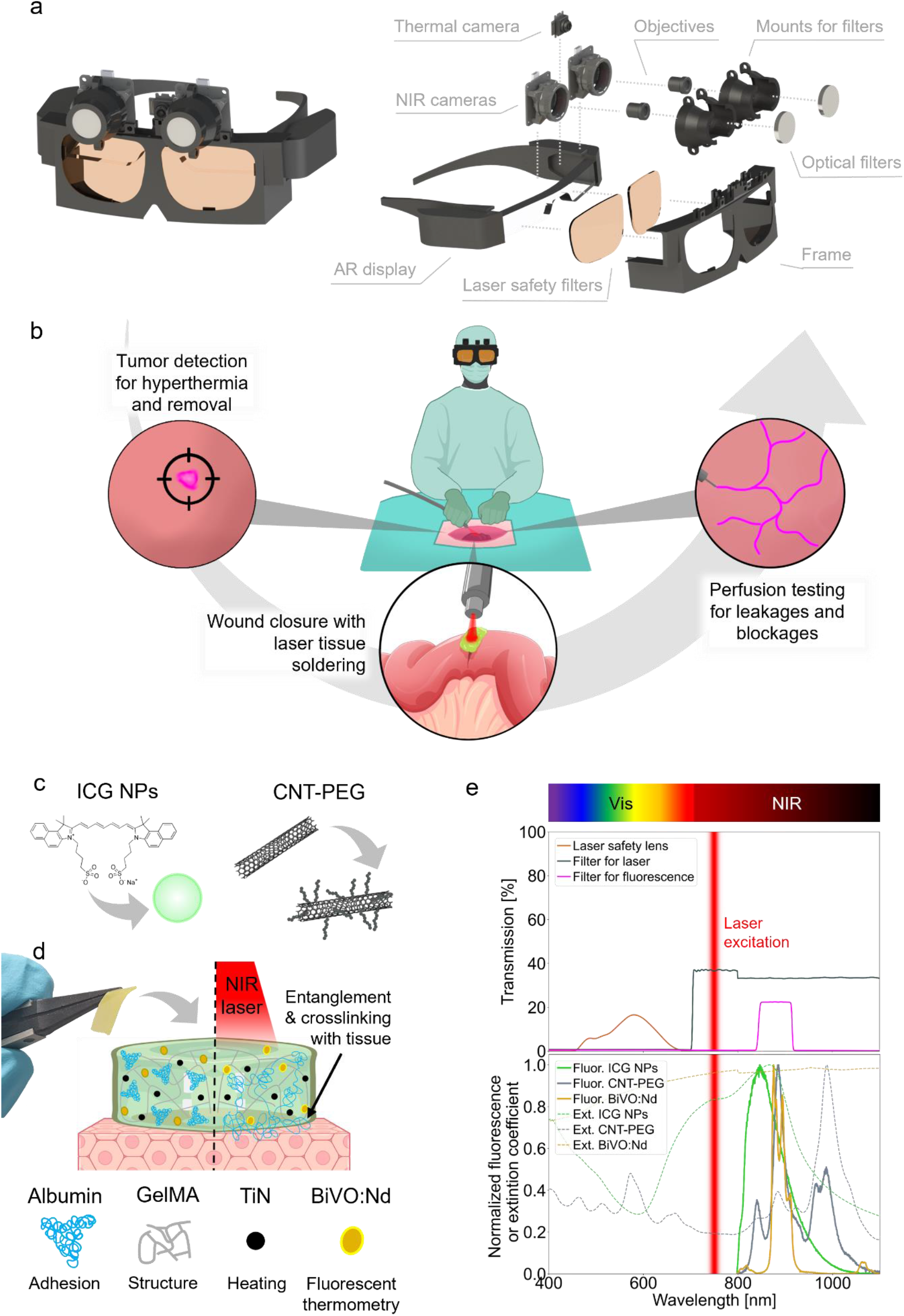
design of the augmented reality visor for laser tissue soldering. (a) Assembled and exploded views of the AR visor with its component. (b) Illustration of the applications of the visor in various surgical procedures: surgical guidance and tumor hyperthermia, wound closer with laser tissue soldering, and vessel perfusion. (e) Illustration of ICG NPs and the molecular structure of ICG, as well as SWCNT and CNT-PEG (sketch for visualization). (d) Illustration of the mechanisms of laser tissue soldering with paste picture and main components. (e) Transmission spectra of the filters used in the AR visor. Absorption and emission spectra of the nanothermometers used for soldering, and ICG NPs and CNT-PEG used as contrast agent.

To ensure user safety from laser exposure, the visor incorporates lenses adapted from commercially available safety goggles. The design of the visor ensures that light entry is restricted exclusively to the lenses, preventing any peripheral light intrusion from the sides of the visor. The frame of the visor is designed to mount onto the smart glasses, equipped with three specialized cameras: one for monitoring laser spot size and position, another for detecting fluorescence, and a third for temperature monitoring during illumination (Figure 1a). NIR cameras using CMOS detectors equipped with NIR-compatible objectives are employed to record the laser light and the fluorescence. Optical filters are used to select the desired wavelengths: a 700 nm longpass (LP) filter is used to record the laser light, while an 880 nm bandpass (BP) filter with a full width at half maximum (FWHM) of 70 nm isolates the fluorescence light, substantially reducing light and laser excitation. The datasets are then overlayed semi-automatically depending on the working distance.

To evaluate the efficacy of the AR visor to assist in fluorescence-guided surgery, we conducted tests using two distinct types of nanoparticles (Figure 1d). The first type included nanoparticles incorporating the medically approved dye indocyanine green (ICG). ICG nanoparticles (ICG NPs) are particularly advantageous due to their enhanced photostability and superior ability to be taken up by tumors, whether through passive or active targeting mechanisms. Specifically, the ICG NPs used in this study were composed of polyallylamine, encapsulated with a dextran coating to improve stability.(*52*) A significant benefit of these particles is their tunable size, which can be precisely adjusted to optimize biodistribution and cellular uptake. Additionally, we examined fluorescent single-walled carbon nanotubes (SWCNT). These nanoparticles can be non-covalently functionalized through adsorption of macromolecules such as phospholipid-polyethylene glycol conjugates (PEG), resulting in PEGylated carbon nanotubes (CNT-PEG). CNT-PEG demonstrates increased stability within saline and blood environments compared to non-functionalized SWCNTs, maintaining their fluorescence properties effectively under these conditions.(*53*) Both types of nanoparticles can be efficiently excited by the 750 nm laser utilized in our AR visor system, and they emit fluorescence within the wavelength range detectable by the fluorescence camera (Figure 1e). This capability allows for precise imaging and targeting during surgical procedures, underscoring the potential of using varied nanomaterials in enhancing the accuracy and outcomes of fluorescence-guided surgeries.

For laser tissue soldering in wound closure, we employed an nanoparticle-enhanced solder paste (Figure 1c).(*54, 55*) This paste features albumin for tissue adhesion via thermal denaturation, crosslinked methacryloyl gelatine for ease of shaping and application, and titanium nitride (TiN) nanoparticles to enhance photothermal efficiency. The use of TiN nanoparticles allows for lower, safer laser powers while achieving the same thermal effect. Additionally, the paste incorporates fluorescent bismuth vanadate nanoparticles doped with neodymium (BiVO_4_:Nd^3+^, here referred to as BiVO:Nd), serving as fluorescent nanothermometers to guide the procedure.(*56*)

### Information visualization and tumor detection with AR guidance

To enhance the precision of feature detection, particularly tumor identification, our system integrates simultaneous visualization of excitation areas and areas emitting strong fluorescent signals. Additionally, temperature visualization assists in monitoring whether therapeutic hyperthermia is achieved or if unwanted heating occurs. This multimodal information is represented using distinct color codes to facilitate intuitive interpretation by surgeons. This information is displayed through a compact visor as shown in Figure 2a and the Supporting Information (Table S1).

**Figure 2:**
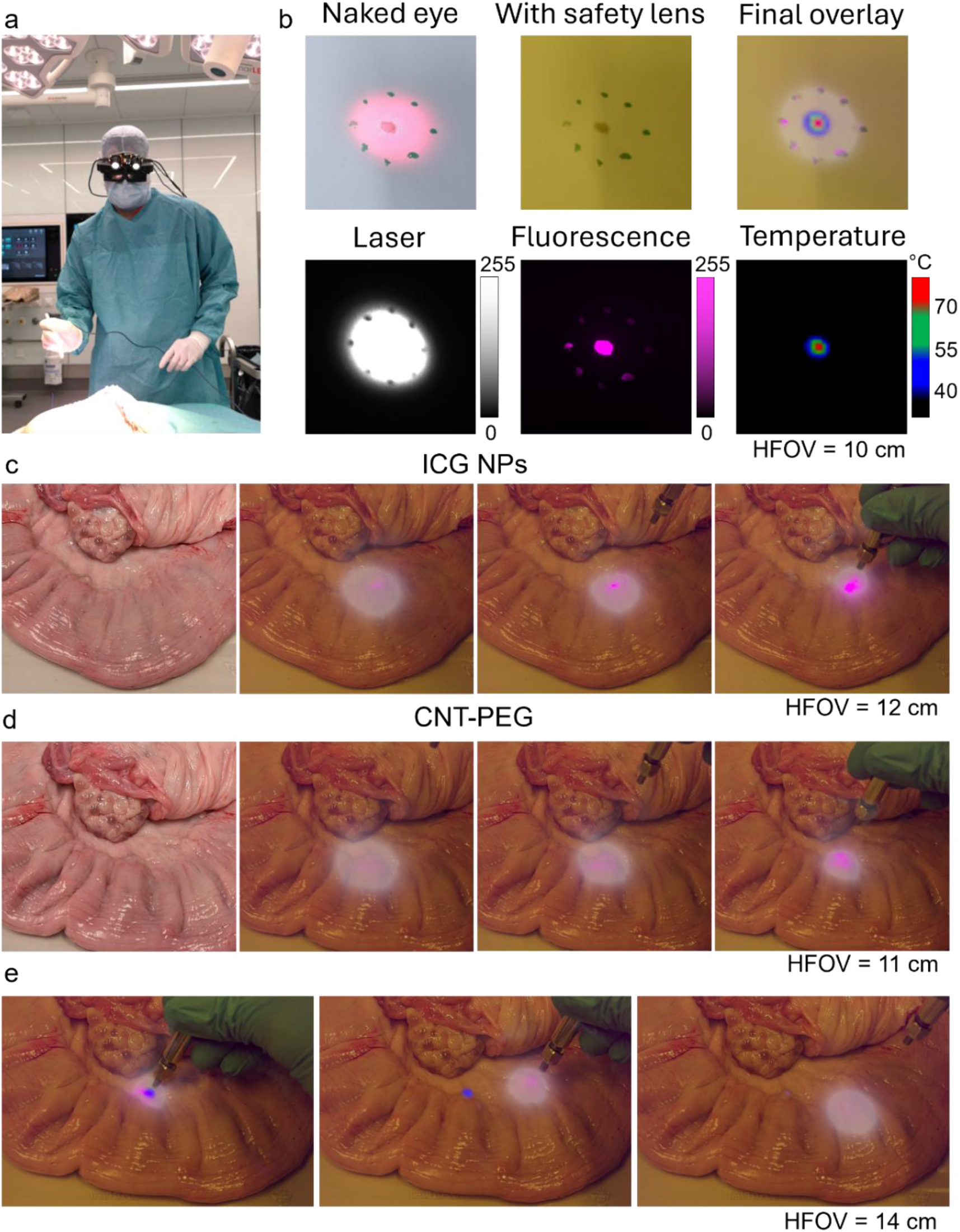
Information display and tumor detection for removal guidance and hyperthermia. (a) Photo of the visor being worn by a surgeon in the OR. (b) Example of how the different images are overlayed on the naked eye field of view using drops of ICG as fluorescent sample (6 *µ*l central dot, 3 *µ*l surrounding dots). Horizontal field of view (HFOV) of about 10 cm. (c) Detection of area with ICG NPs injected into bovine uterus. Naked eye vision (first image on the left) cannot distinguish which areas are labelled and which are not. The AR visor can aid by visualizing the fluorescence which can be seen at various laser illumination distances between 1 and 10 cm (three images on the right). (d) Detection of labelled area in bovine uterus can also be seen using CNT-PEG as labelling agents. (e) Hyperthermia (heating of tissue by the laser) can be performed in this case, as seen from the images taken at different time steps. The color-coded heatmap allows the surgeon to avoid reaching temperatures that are too high by moving the laser away from the region and waiting for it to cool down, as seen from disappearing blue area in the second and third panel.

The 750 nm laser used in our system is partially visible to the human eye, despite the declining sensitivity of both the human visual system and silicon-based detectors to NIR wavelengths beyond 700 nm. This phenomenon is illustrated in Figure 2b, where drops of ICG on a surface can be faintly seen. Importantly, the perceived brightness of NIR light underestimates its actual intensity, presenting a potential safety hazard. To mitigate this risk, we employ laser-safe lenses that block the laser light effectively. However, these lenses critically alter the perception of other colors due to the absorption of various visible wavelengths, necessitating the pre-lens display of information to utilize the full color spectrum. In our visual interface, the excitation source appears as a white halo, while areas of fluorescence are depicted in magenta to contrast sharply against the yellow tint produced by the laser-safe lenses (Figure 2b). To avoid overwhelming the user with data, we employ a simplified, color-coded temperature map. Temperatures below 40 °C are not shown; those slightly above normal body temperature yet below 55 °C, which could cause unintended tissue damage on the surrounding tissue, are shown in blue. Temperatures optimal for laser tissue soldering, between 55 °C and 70 °C, are shown in green, while potentially harmful temperatures above 70 °C are indicated in red. The images shown here were captured by a camera, and the overlays were artificially recreated for demonstrative purposes.

To provide surgical guidance and possible heating capabilities, there is a need to use fluorescent agents. For experimental validation, we simulated targeted features by injecting 0.1 ml of saline solutions containing either ICG NPs (OD 1.35) or CNT-PEG (OD 17) into bovine mesometrium (Figure 1b-d). These areas, invisible to the naked eye, emit discernible fluorescence signals when viewed through the AR visor, even under broad illumination. The fluorescence from ICG NPs is notably stronger than that from CNT-PEG, which has a lower excitation efficiency at 750 nm and emits more strongly around 950-1000 nm, nearing the detection limit of silicon-based detectors. Despite not being specifically optimized for CNTs, their fluorescence is effectively captured by our system, especially at closer ranges, underscoring the versatility and efficacy of our AR visor in fundamental research and clinical applications. An additional advantage of the system is that the surgeon can move the excitation source across the area to identify other labeled regions or closer to an identified area to enhance the signal clarity. The thermal camera aids in monitoring for unintentional warming during visualization or intentional heating in tumor treatments, improving the accuracy of therapeutic interventions. Color coding the thermal output also facilitates achieving specific therapeutic temperature ranges, which is currently challenging with systems being used in the clinic. Typically, temperatures around 45 °C are desirable for therapeutic effects, with higher temperatures potentially leading to adverse effects. Thus, areas appearing blue are within the desired range for hyperthermia, with the process paused when areas turn green, as demonstrated in experiments with CNT-PEG labeled regions (Figure 1d).

### Laser tissue soldering with augmented visor

Following tumor resection, wound closure is often a critical step. Laser tissue soldering offers an alternative to sutures for closing wounds in soft tissues. Effective laser tissue soldering requires meticulous temperature control to ensure proper bonding. The desired bonding temperature, between 55 and 70 °C, is color-coded green on the display of the visor. Temperatures below this threshold that do not facilitate bonding appear blue, while potentially harmful temperatures that could cause excessive thermal damage are displayed as red.

The AR visor has been tested by using it to perform soldering on an ex vivo porcine blood vessel. A piece of solder paste (5×3 mm) was placed on top of the vessel and then soldered on top of it using the 750 nm laser. The display of the laser illumination helps with guiding the laser to the soldered area, while fluorescence of the paste also helps with visualizing the position of the paste, which could be less visible due to the presence of the safety filters. Additionally, temperature display provides further guidance. When the temperature is rising too fast, the laser can be moved away while the temperature at the solder location can still be monitored while it cools down (Figure 3a).

**Figure 3:**
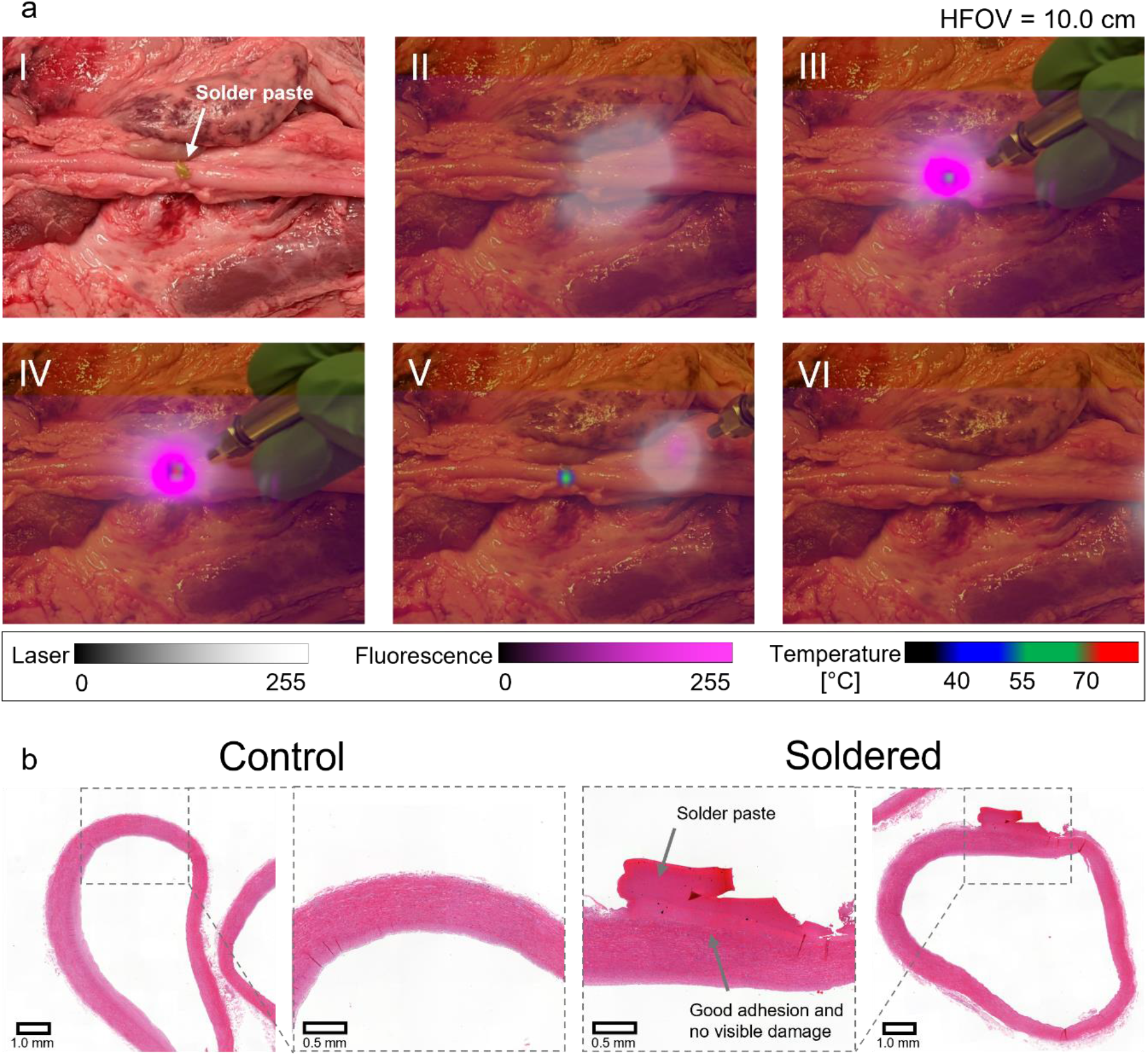
Soldering using the AR visor. (a) Laser tissue soldering of a porcine blood vessel guided by the AR visor at various time steps. I) Naked-eye vision of the vessel with solder paste on it. II) Laser hovered over the region to better locate the paste. III) Paste localized and heating started. IV) In situations where temperatures exceed the required levels, they are indicated in red, alerting the user to act. V) The laser is moved away from the paste to avoid overheating, while the temperature of the paste is monitored during cooling. VI) The paste has returned to its initial temperature. (b) H&E-stained histological slides showing a healthy native blood vessel (left), compared with soldering with the AR visor (right). Good adhesion of the paste on the tissue can be noticed, as well as no visible thermal damage can be seen in the soldered sample.

The panels in Figure 3a illustrate the various stages of the soldering process, including initial paste location confirmation (II) and continuous heating (III). If excessive temperatures are detected (IV), the laser is moved away from the target area, and the cooling process is monitored (V, VI). This method allows for precise temperature control to maintain it at the optimal level. The paste emits intense fluorescence when illuminated from a close distance, creating a magenta-shaded halo without the distinct border of the paste itself. The size of the halo is directly related to the distance of the laser and its positioning. This feature can assist the user in fine-tuning their positioning during the procedure.

Histological analysis of a soldered porcine blood vessel confirms the efficacy of the AR visor-guided laser tissue soldering (Figure 3b). The results demonstrate excellent adhesion and fusion between the tissue and the solder paste, with no visible thermal damage compared to control samples. This evidence underscores the AR visor’s potential to enhance surgical outcomes through precise, controlled application of emerging laser soldering technology, ensuring effective and damage-free tissue bonding.

### Visualization of blood perfusion

In the evaluation of the AR visor’s effectiveness in organ perfusion assessment, we sought to demonstrate a significant enhancement in the visualization of vascular structures during perfusion tests on bovine mesometrium. Such structures are not readily discernible to the naked eye (Figure 4a, top), even after injecting blood labelled with ICG NPs (OD 0.27). However, with the use of the AR visor, the detailed architecture of the blood vessel tree becomes distinctly visible. As the blood is injected, it is possible to track its flow through the vessel structure by accordingly moving the illumination fiber. Optimal visibility of the entire structure is achieved by maintaining the fiber at a distance of approximately 30 cm, adequately illuminating the area. This marked improvement highlights the visor’s capability to detect subtle anatomical features essential for precision in surgical procedures.

**Figure 4:**
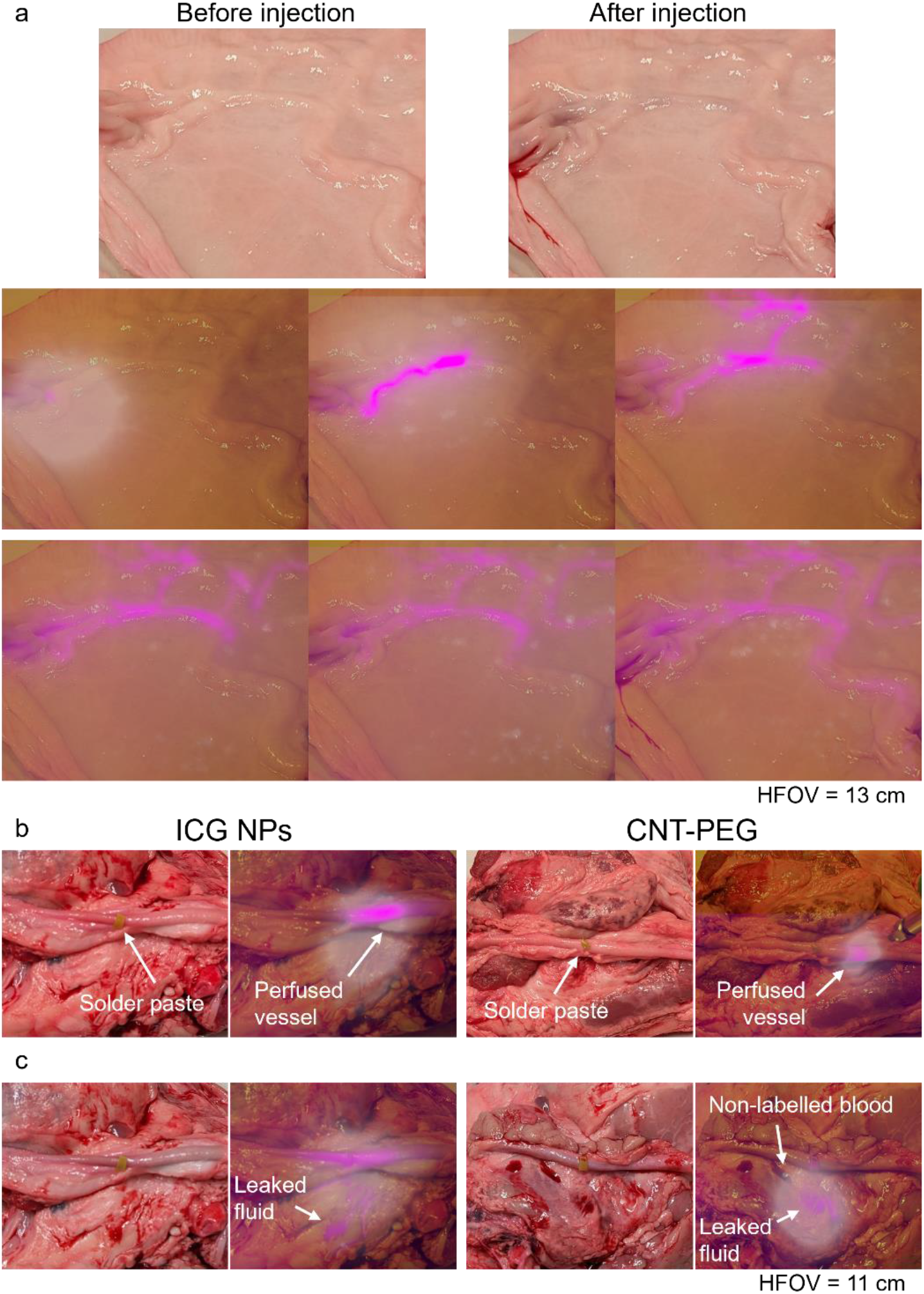
Perfusion testing enabled by the AR visor. (a) Perfusion test performed on a piece of bovine mesometrium. The shape and connectedness of the vasculature is not clearly visible by naked eye evaluation, both before and after injection (top photos). The structure of the blood vessel tree is clearly visible using the AR visor, as the injected blood with ICG NPs spreads in the vessels. (b) Perfusion test on a porcine blood vessel after wound closure using blood labeled with ICG NPs (left side) or a saline solution of CNT-PEG (right side). (c) A simulated leak can be easily picked up by the AR visor, clearly distinguishing labelled blood (ICG NPs on the left, CNT-PEG saline solution on the right) with surrounding blood present due to the surgery.

Further analysis was conducted to validate the efficacy of the system for post-wound closure using two different contrast agents: blood labeled with ICG NPs (OD 0.27) and a saline solution containing CNT-PEG (OD 17). The ICG nanoparticles exhibited strong visibility within the vascular system, providing a clear pattern of blood flow (Figure 4b, left). The visualization with CNT-PEG was less pronounced, as previously mentioned, but still rendered the structure visible (Figure 4b, right). The AR visor also proved highly effective in leak detection during simulated conditions. A leakage scenario was created by placing the labeled fluid adjacent to the vessel, along with unlabeled blood, to replicate a more complex and challenging scenario. The visor successfully differentiated between areas with labeled blood and the surrounding blood, an essential function for identifying and addressing potential leaks during surgery (Figure 4c). This capability is particularly valuable as it helps reduce the risk of postoperative complications by enabling prompt corrective measures. It should also be noted that in these tests, no significant heating was detected by the thermal camera. However, should there be any increase in temperature, it would be displayed, allowing the user to take appropriate actions.

Overall, the integration of the AR visor into surgical practice offers substantial improvements in the precision and safety of surgical interventions. The enhanced visual capabilities provided by the AR visor ensure that surgeons can perform delicate procedures with greater accuracy and confidence. Depending on the specific application, the use of ICG nanoparticles can be more advantageous with the current setup due to their superior visualization properties. However, CNT-PEG nanoparticles still remain a viable option for certain scenarios, allowing for flexibility in choosing the appropriate imaging agent to meet the unique requirements of each surgical procedure. Based on the modular approach of the AR visor, this technique could be rapidly applied to other fluorescence-guided imaging agents, hence improving the availability of this surgical method, which could ultimately advance patient care.(*57*) This technology facilitates a significant advancement in medical imaging, optimizing the visualization necessary for complex surgical tasks and enhancing the overall efficacy of medical procedures.

## CONCLUSIONS

In this study, we have successfully demonstrated significant advancements in surgical visualization and intervention made possible by the integration of an AR visor. The AR visor, equipped with specific imaging enhancements, has proven to be a valuable tool in various simulated surgical scenarios, enhancing the precision and safety of procedures. The visualization capabilities of the AR visor were extensively demonstrated in perfusion tests on porcine and bovine tissues and in the application of laser tissue soldering on ex vivo porcine tissues. The use of ICG NPs in these tests showed distinct advantages in terms of visibility and integration within the vascular system, enabling surgeons to perform with greater accuracy. Furthermore, the AR visor’s ability to differentiate between labeled and surrounding blood during leak detection scenarios has underscored its utility in preventing postoperative complications and ensuring successful surgical outcomes. Moreover, by evaluating CNT-PEG, we demonstrate that other promising NIR fluorescent dyes and nanoparticles, for which this system is not fully optimized, can still be effectively detected by the visor and fulfill their intended use. This underscores the AR visor’s capability to adapt to a range of fluorescent agents, enhancing its versatility across different surgical applications.

Considering possible future developments, there are some potential enhancements that could further improve the AR visor’s application in surgical settings. Developing algorithms for automatic adjustment of image overlays based on real-time data could provide even faster and more precise overlays, thereby decreasing procedure times and enhancing outcome predictability. Additionally, expanding the range of detectable fluorescence ranges could allow for the simultaneous use of multiple fluorescent agents, broadening the scope of surgical applications, or of detection of different emission peaks, enabling the measurement of temperature through ratiometric fluorescent thermometry.

In conclusion, the AR visor represents a transformative advancement in medical technology, offering enhanced visualization that not only translates into safer, more precise surgical interventions with established techniques but also facilitates the integration and adoption of various emerging technologies. Its development and refinement continue to push the boundaries of what is possible in surgical care, promoting greater improvements in patient outcomes and encouraging the surgical community to more readily adopt these promising new technologies as the technology evolves.

## EXPERIMENTAL

### Augmented reality glasses

A Moverio BT-40 (Epson) was used as augmented reality visor. The attachment for the goggles and the various add-ons (cameras, laser safety lenses) was 3D printed (Prusa i3 MK3S+) with PETG black filament. The lenses of commercially available laser safety goggles (Thorlabs, LG9) were removed from the original frame and inserted in the attachment. To record the laser light and the fluorescence two NIR cameras were used (U3-3680XLE-NIR-GL 1/2.5” NIR USB3 Camera, IDS Imaging). Each camera had a NIR lens screwed in front of the sensor (f/2.5, NIR, 8.0mm HEO Series M12 Lens, Edmund Optics). In front of the camera for the laser detection (on the left side of the visor) a 700 nm longpass filter (Thorlabs, FEHL0700) was used. In front of the camera used to record the fluorescence an 880 bandpass filter (Thorlabs, FBH880-70) was used. The filters were held using 3D printed parts. The gains and exposure time were calibrated to achieve higher signal quality without increasing the time delay. A thermal camera (Tiny1-C 04312X l, InfiRay) with 25 Hz frame rate, 256×192 resolution, and 40°×30° FOV was used to record the thermal images. Further information can be found in the supplementary information.

High speed cables were used to connect the cameras and the visor to a small portable computer (MOREFINE M6 Mini PC, Intel 11th Gen N6000) which could be either worn by the user or placed nearby. A code in Rust was used to process the data from the cameras and to display the output on the visor. The code also allowed to manually align the images to the field of view by using the keyboard: letters were used to select the image to align, arrows to move each image, and “+” and “-” to scale the figure.

The output of the cameras was color-coded. For the excitation source the color white was used, for the fluorescence the color magenta was used. The thermal image was color coded using three colors as base: blue for temperatures in the range 40-55 °C; green for temperatures in the range 55-70 °C; red for temperatures above 70 °C. A transition transient of 5 °C was used before the start of each range by linearly change the RGB values of the previous range to the next one. In order to have better visualization the intensity coming from the excitation image was set to 50% of the other images. The displayed color for pixels with information coming from more than one camera was displayed using the following order of priority: temperature, fluorescence, excitation.

Overlays in the figures were recreated artificially for demonstrative purposes. The headset was placed at an angle of 35° and distance to the sample was set at about 35 cm with the goal of mimicking its use during surgery. A smartphone (Google Pixel Pro 8) was used to take pictures and placed right in front of the visor to mimic the same field of view of a person wearing the visor. A laser safety lens was placed in front of the camera for the pictures where the overlay was done. The white balance value was kept constant, using the value of the picture without the laser safety filter on.

### Laser system

As excitation and heating source, a 750 nm CW laser (FC-750/5W, CNI Lasers) with a maximum output power of 5 W was used. A portable optical module was used to increase the monochromacity of the laser light and avoid being picked up by the fluorescence camera. This module included an optical fiber (NA 0.22, Ø400 μm) to guide the laser from the laser head to the optical module. a collimating plano-convex lens (f = 30 mm, 780 nm V-coating; Thorlabs), a 750 nm bandpass filter (FWHM = 40 nm; FB750-40, Thorlabs), and a plano-convex lens (f = 30 mm, 780 nm V-coating; Thorlabs) for coupling with the output fiber. An optical fiber with NA 0.22 and Ø400 μm was used to shine the laser on the samples. After passing the optical module and the medical fiber the maximum available power was reduced to 2.5 W.

### Solder paste formulation

The solder paste was prepared as previously described.(*54, 55*) Briefly, water solutions of bovine serum albumin (BSA, Sigma–Aldrich, A2153), gelatin modified with methacryloyl (GelMA), 0.0069% LAP (Lithium phenyl-2,4,6-trimethylbenzoylphosphinate, Sigma–Aldrich, 900889), flame-made BiVO_4_:Nd^3+^ nanoparticles, and TiN nanoparticles (PlasmaChem GmbH, PL-HK-TiN) were quickly mixed using a vortex and then casted into the appropriate mold. The paste was cooled for at least 1 h at 4°C and then crosslinked under UV irradiation for 5 min (UVASPOT 400T, Dr. Hönle AG). BiVO nanoparticles were prepared as described previously.(*54*) and then crosslinked under UV irradiation for 5 min (UVASPOT 400T, Dr. Hönle AG).

### ICG nanoparticles synthesis

The synthesis of the ICG nanoparticles was conducted adapting the protocol by Yaseen et al.(*52*) A 3 ml solution of 0.05 M Disodium Phosphate (Sigma Aldrich) and MilliQ water was lightly mixed with a pre-cooled 0.5 ml solution of 65 kDa polyallylamine (PA, Sigma Aldrich) and HCl (Sigma Aldrich) solution at 4 °C. The solution was made by mixing 1.94 ml of water with 31.6 *µ*l of HCl and 20 *µ*l of PA, with a final concentration of 1 mg/ml of PA. Mixing was done using a vortex for 15 s in a 15 ml centrifuge tube. Then, 3 ml of 1 mg/ml ICG (Carl Roth, 7695.2) solution was mixed and set aside at 4 °C for 10 min. Afterwards, 1.5 ml of 2 mg/ml dextran solution (70 kDa, Sigma-Aldrich, 31390-25G) was added, mixed using a vortex for 30 s and then aged for 2 h at 4 °C. The sample was centrifugated at 12800 × g for 30 min and resuspended in either MilliQ water or phosphate buffered saline (PBS). The centrifugation process was repeated one more time. The concentration of ICG NPs was measured by absorbance spectroscopy and hemocytometry. An OD of 1.35 is equivalent to 2.2·10^8^ particles/ml of ICG in the solution.

### CNT preparation

SWCNTs were modified with PEG-PL as described by Nissler et al.(*17*) In short, 2 mg/ml (6,5)-Chirality-enriched CoMoCat SWCNTs (Sigma-Aldrich, 773735) were dispersed (10 min 90% amplitude-Sonics Vibra cell VCX 500, cup horn) in sodium deoxycholate (DOC, 1% w/w in H_2_O), followed by centrifugation (20 min at 7800 × g) to remove non-dispersed SWCNTs. 4 ml of concentrated DOC-SWCNTs were transferred into a centrifugal filter (Amicon Ultra-4, PLHK Ultracel-PL Membran, 100 kDa) to exchange the surface modification to sodium cholate (SC, 12 mg/ml) by repeated spin filtration (3×5 min at 5000 × g). 4 ml SC-SWCNTs were mixed with 10 mg PEG-PL (18:0 PEG5000 PE, 1,2-distearoyl-sn-glycero-3-phosphoethanolamine-N-[methoxy(polyethylene glycol)-5000], Avanti Polar Lipids) dissolved in 1 ml 1 × PBS and dialyzed 4 days against 1 × PBS, using a 1 kDa cutoff dialysis tube (Spectra/ Por®). The concentration of PEG-PL-SWCNTs was measured by absorbance spectroscopy.(*18*) An OD of 17 is equivalent to 67 *µ*g/ml.

### Materials characterization

The measurement of the extinction properties of the various materials were made using a UV-Vis NIR spectrophotometer (Cary 5000 UV-Vis-NIR Spectrophotometer, Agilent), using a step of 1 nm and integration time of 0.1 s. Suitable blank and zero references were also recorded and used to automatically calculate the absorbance or transmission of the samples. Fluorescence spectra was acquired using a NIR spectrometer (100 μm slit; STS-NIR, Ocean Optics) with the previously mentioned laser as excitation source. Particle concentration was measured through hemocytometry. Further characterization is present in the supplementary information.

### Tumor visualization model

A bovine uterus was retrieved from the local slaughterhouse on the same day as the experiment. 0.1 *µ*l of a solution of ICG NPs (1.35) or of PEG-CNT (OD 17) was injected in a randomly selected region of the uterus to simulate a region where the nanoparticles were able to target a tumor region. Then, the laser was shined at the specimen and images taken at various times during the application. The laser was moved in order to find the labeled area, check surround areas, and heat the specimen. The experiment was conducted both at room temperature and with the tissue warmed by hot water to around 37 °C.

### Laser Tissue Soldering

Porcine carotid arteries and jugular veins were retrieved from the local slaughterhouse on the same day of the experiment. Some additional porcine tissue (heart or other muscular tissue) was used as background for the experiment in order to have a more realistic setting. A piece of solder paste (5×3 mm) was placed on top of the tissue. Then, the laser at was shined on the tissue for around 1 min making sure not to overshoot the temperature, apart from the cases where it was done on purpose for demonstrative purposes. Images were taken at various times during the application.

Soldering was also performed on blood vessels of different size which were then collected for histological analysis, together with reference samples. Samples were placed in a formalin solution (Histofix 4%, Carl Roth, P087.5) and processed by an external service (SophistoLab AG). The histological slices were images using a slide scanner at ×20 magnification (Slide Scanner Pannoramic 250, 3D Histech).

### Blood vessel tree visualization and perfusion test

Porcine blood with anticoagulant was taken from the local slaughterhouse and used on the same day. A solution of ICG NPs (OD 1.35) was added to the blood in a ratio of 1:5, leading to a concentration of 4.5·10^7^ particles/ml ICG NPs in blood. Around 0.5 ml were injected in the blood vessels of bovine mesometrium while illuminated by the laser. Pictures were taken during the injection while the laser was used to see the flow of the blood in the vessel tree.

For the perfusion test, porcine carotid arteries and jugular veins together with additional porcine tissue (heart or other muscular tissue) used as background were used. The perfusion was carried our either using a 1:5 solution of ICG NPs in blood or a CNT-PEG saline solution (OD 17). Pictures were taken during the injection. Moreover, a leaking sample was simulated by adding the solutions on the tissue nearby. Additional blood was added on the tissue in the vicinity of the vessel to simulate a more challenging scenario.

## Acknowledgments

We acknowledge funding from the Swiss National Science Foundation (Eccellenza grant no. 181290) and the Oertli Foundation. The authors gratefully acknowledge ScopeM for their support & assistance in this work. Some illustrations were created from illustrations from BioRender.

