## Supplementary information for "An Augmented Reality Visor for Intraoperative Visualization, Guidance and Temperature Monitoring using Fluorescence"

### Augmented Reality Visor for IR- and NIR-Guided Laser Tissue Soldering Integrated in Surgical Workflows

### Nanoparticle characterization

DLS measurement (Litesizer 500, Anton Paar) was used to measure the hydrodynamic size of the particles (Figure S1a).

Electron microscopy images confirm the findings from the DLS measures. SEM images were carried out using a scanning electron microscope (Magellan, Thermo Fisher Scientific) from ScopeM. The machine was operated using an acceleration voltage of 10 kV, beam current of 0.20 nA and a working distance of 10 mm. The following images show the structure of the ICG NPs after being dropcasted on a Si wafer (Figure S1b).

a

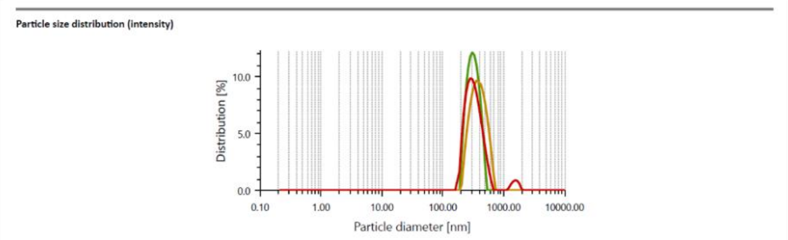


Measurements - Number

| Index | Color | Hydrodyn. diam. [nm] | Polydisp. index [%] | Peak 1 [nm] | Peak 2 [nm] | Peak 3 [nm] | Transmittance [%] |
|---|---|---|---|---|---|---|---|
| 1 | green | 355.2 | 21.7 | 225.5 (Number) | 335.8 (Number) | - (Number) | 73.2 |
| 2 | orange | 430.5 | 9.5 | 363.2 (Number) | - (Number) | - (Number) | 74.2 |
| 3 | red | 387.3 | 9.3 | 297.4 (Number) | - (Number) | - (Number) | 75.8 |

b

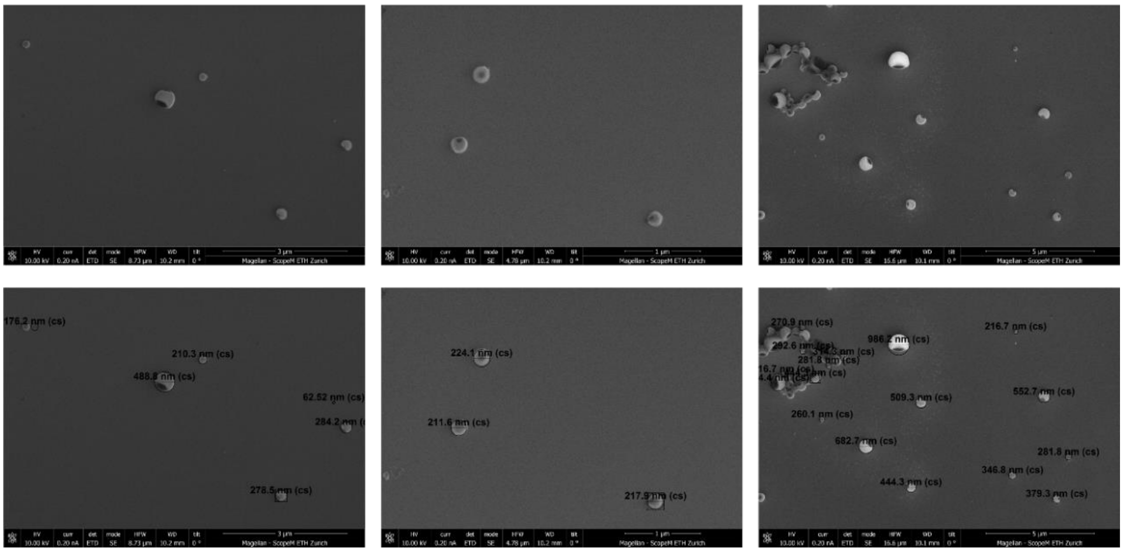

#### Weight and cost

An analysis of the weight of the headset allows to understand how much the setup will disturb the surgeon. Another important parameter is represented by the cost of the visor, which could hinder widespread adoption, especially in developing countries, if too high. The following table shows a breakdown of weight and cost for each component. (Table S1)

Table S1: breakdown of the approximate weight and cost of the various components of the headset.

| **Item** | **Weight [g]** | **Cost [€]** |
|---|---|---|
| AR visor (excl. cables and controller) | 70 | 700 |
| 3D printed add-on | 20 | Negligible |
| Laser safe lenses | 10 | 200 |
| NIR camera (x2) | 20 (x2) | 300 (x2) |
| Thermal camera | 5 | 240 |
| Optical filter: LP 700 | 2.5 | 130 |
| Optical filter: BP 880 | 2.5 | 130 |
| **Total headset** | **150** | **2000** |
| Laser setup (including accessories) | Not relevant | 4000-5000 |
